# Exercise Mitigates Flow Recirculation and Activates Mechanosensitive Transcriptome to Uncover Endothelial SCD1-Catalyzed Anti-Inflammatory Metabolites

**DOI:** 10.1101/2023.05.02.539172

**Authors:** Susana Cavallero, Mehrdad Roustaei, Sandro Satta, Jae Min Cho, Henry Phan, Kyung In Baek, Ana M. Blázquez-Medela, Sheila Gonzalez-Ramos, Khoa Vu, Seul-Ki Park, Tomohiro Yokota, Jennifer A. Sumner, Julia J. Mack, Curt D. Sigmund, Srinivasa T. Reddy, Rongsong Li, Tzung K. Hsiai

## Abstract

Exercise modulates vascular plasticity in multiple organ systems; however, the metabolomic transducers underlying exercise and vascular protection in the disturbed flow-prone vasculature remain under-investigated. We simulated exercise-augmented pulsatile shear stress (PSS) to mitigate flow recirculation in the lesser curvature of the aortic arch. When human aortic endothelial cells (HAECs) were subjected to PSS (*τ*_ave_ = 50 dyne·cm^−2^, ∂τ/∂t = 71 dyne·cm^−2^·s^−1^, 1 Hz), untargeted metabolomic analysis revealed that Stearoyl-CoA Desaturase (SCD1) in the endoplasmic reticulum (ER) catalyzed the fatty acid metabolite, oleic acid (OA), to mitigate inflammatory mediators. Following 24 hours of exercise, wild-type C57BL/6J mice developed elevated SCD1-catalyzed lipid metabolites in the plasma, including OA and palmitoleic acid (PA). Exercise over a 2-week period increased endothelial SCD1 in the ER. Exercise further modulated the time-averaged wall shear stress (TAWSS or *τ*_ave)_ and oscillatory shear index (OSI_ave_), upregulated *Scd1* and attenuated VCAM1 expression in the disturbed flow-prone aortic arch in *Ldlr*^-/-^ mice on high-fat diet but not in *Ldlr*^-/-^*Scd1*^EC-/-^ mice. *Scd1* overexpression via recombinant adenovirus also mitigated ER stress. Single cell transcriptomic analysis of the mouse aorta revealed interconnection of *Scd1* with mechanosensitive genes, namely *Irs2*, *Acox1* and *Adipor2* that modulate lipid metabolism pathways. Taken together, exercise modulates PSS (*τ*_ave_ and OSI_ave_) to activate SCD1 as a metabolomic transducer to ameliorate inflammation in the disturbed flow-prone vasculature.

## INTRODUCTION

Physical exercise augments blood flow in the cardiovascular system and modulates molecular transducers to ameliorate metabolic disorders ^1^. During exercise, increased cardiac contractility and heart rate enhance PSS to improve endothelial cell plasticity ^2^, to mitigate atherosclerosis ^3, 4^, and to delay cognitive aging and neurodegeneration ^5–8^. PSS is unidirectional and axially aligned with the flow direction in the straight regions of the vasculature, and PSS is known to activate antioxidants, including superoxide dismutase (SOD), and to attenuate proinflammatory cytokines, adhesion molecules, and NADPH oxidase system ^9–12^. In contrast, disturbed flow, including oscillatory shear stress (OSS), is characterized as bidirectional and axially misaligned at the bifurcating regions or the lesser curvature of the aortic arch. OSS is well-recognized to induce oxidative stress and inflammatory response ^9, 13–15^. However, the specific exercise-activated metabolomic transducers that mitigate inflammation and atherosclerosis in the disturbed flow-prone vasculature are not completely elucidated ^16^.

Several mechano-sensitive molecular transducers have been reported to regulate fatty acid (FA) uptake and transport in the vascular endothelium, including vascular endothelial growth factor B (VEGFB) ^17, 18^ and peroxisome proliferator activated receptor gamma (PPARγ) ^19^. PPARγ is a member of the nuclear receptor superfamily, participating in the regulation of blood pressure, hypertension, hyperlipidemia, and insulin sensitivity ^20–22^. While laminar shear stress upregulates SCD1 in human aortic endothelium through a PPARγ-dependent mechanism *in vitro* ^23^, we investigated whether physical exercise activates PPARγ-SCD1 pathway to promote vascular protective metabolites in the disturbed flow-exposed aortic arch.

SCD1 is a key transmembrane enzyme in the ER, catalyzing the conversion of saturated long-chain fatty acids (SFA) to Δ9-monounsaturated fatty acids (MUFA). The principal metabolite of SCD1 is oleic acid, which is formed by desaturation of stearic acid. The ratio of stearic acid to oleic acid is implicated in the regulation of cell growth and differentiation through its effect on cell membrane fluidity and signal transduction ^24^. In this context, we performed *in silico* analysis to simulate exercise-augmented PSS to mitigate flow recirculation or disturbed flow in the lesser curvature of aortic arch. Our unbiased and untargeted metabolomic analyses uncovered that PSS activates endothelial SCD1 to catalyze the production of OA as a protective lipid metabolite, as evidenced by the attenuation of NF-κB-mediated inflammatory mediators and ER stress. To demonstrate SCD1 activation in the disturbed flow-prone aortic arch, we further developed mice with endothelial-specific SCD1 deletion (*Ldlr* ^-/-^ *Scd1* ^EC-/-^) to undergo supervised exercise in a voluntary wheel running system. We demonstrated that exercise modulates PSS (*τ*_ave_ and OSI_ave_) and activates endothelial SCD1 as a metabolomic transducer to catalyze lipid metabolites that ameliorate inflammation and atherosclerosis in the disturbed flow-exposed vasculature.

## RESULTS

### Pulsatile shear stress (PSS) up-regulates endothelial SCD1 to catalyze lipid metabolites

Following well-defined PSS (time-averaged shear stress, *τ*_ave_ = 50 dyne·cm^−2^, and slew rate, ∂τ/∂t = 71 dyne·cm^−2^·s^−1^, at 1 Hz) to HAECs monolayers for 4 hours (**Figure 1A**), untargeted metabolomic analysis demonstrated that the SCD1-catalyzed metabolite oleic acid (a monounsaturated omega-9 fatty acid) was significantly increased (\**p* < 0.05, PSS vs. static condition, n=4) (**Figure 1B and 1E**). However, PSS did not increase the saturated fatty acid precursors palmitic or stearic acid (**Figure 1B**). PSS also significantly increased linoleic acid (a polyunsaturated omega-6 fatty acid) and its derivative azelaic acid. Both metabolite heatmap and Principal Component Analysis (PCA) demonstrated that PSS-mediated metabolites were separated from the basal levels (**Figure 1C and 1D**). PSS also increased the production of glycolytic metabolites, including glucose-6-phosphate and fructose-6 phosphate, as previously reported ^25^. Although ECs primarily consume glucose through anaerobic glycolysis ^26^, they contain the enzymatic machinery for FA oxidation ^27^. Consistently, our bioinformatic analysis of compound networks ^28^ showed interconnections among glycolytic and lipid metabolite nodes in HAECs after PSS exposure (**Figure 1F**).

**Figure 1.**
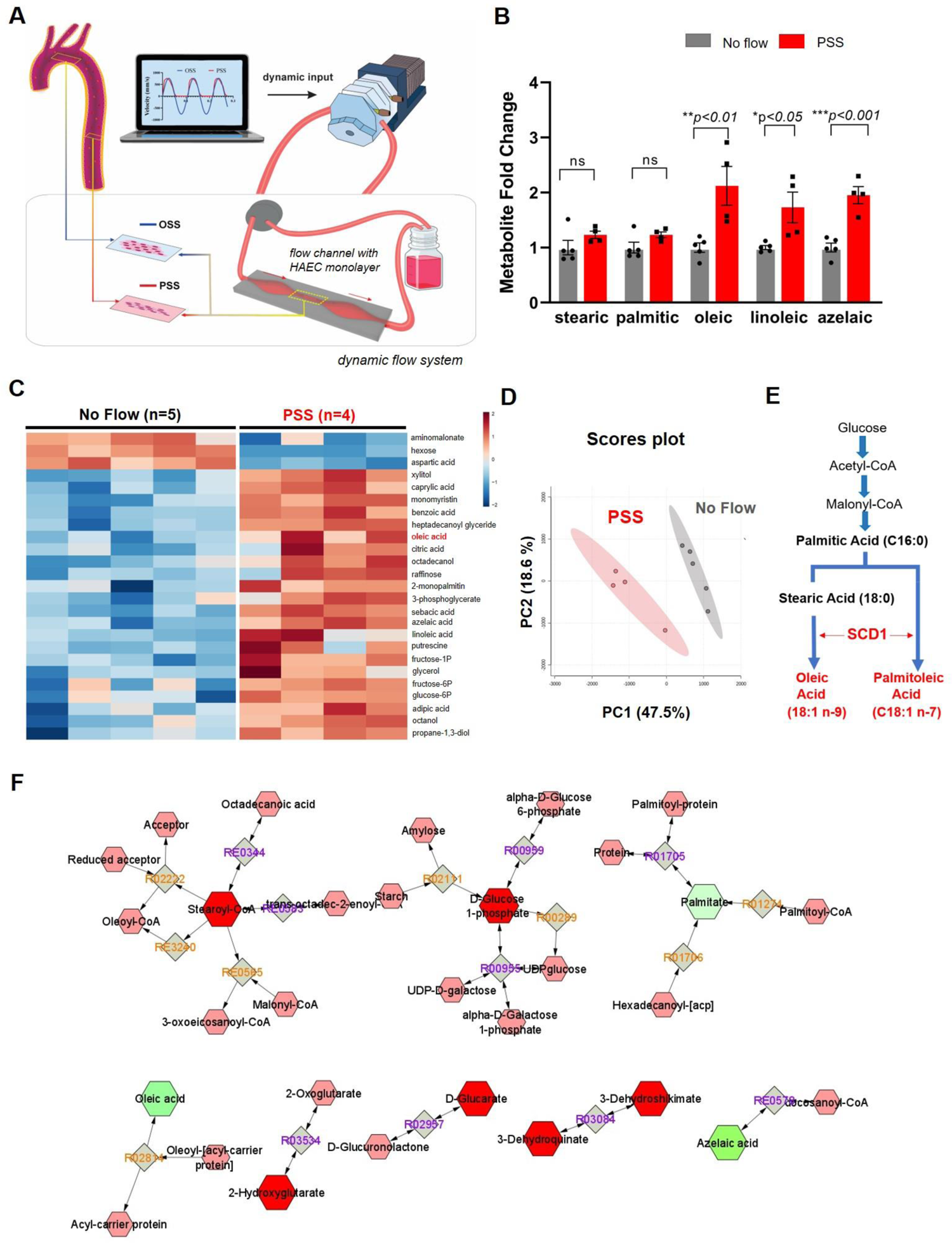
Pulsatile shear stress (PSS) modulates endothelial metabolome and increases SCD1-catalyzed metabolites. **(A)** A custom-built dynamic flow system was used to simulate spatial and temporal variations in shear stress in the arterial system. The machine language (Lab View) drives a peristaltic pump to generate pulsatile wave forms. The configuration of flow channel (contraction and extraction) was designed to provide the well-defined pulsatile flow with the specific slew rates (∂τ/∂t), time-averaged wall shear stress (*τ*_ave_), frequency (Hz), and amplitude to simulate exercise-augmented PSS. The flow channel was maintained inside the incubator at 37°C with 5% CO_2_. A confluent HAEC monolayer was seeded on the glass slides in the flow channel. **(B)** Metabolite samples were collected from HAECs under the static condition (control or no flow, n=5) or PSS (n=4 at 1 Hz for 4 hours) for the untargeted metabolic analysis. PSS significantly increased the lipid metabolites; namely, oleic, linoleic and azelaic acids (\**p* < 0.05 PSS vs. static condition). **(C)** The heatmap reveals an increase in both glycolytic and fatty acid metabolites. The data were analyzed after normalization and scaling using the Pareto method. **(D)** Score plots by the PCA analysis revealed a separation of the representative metabolites. Ellipse represents 90% confidence intervals for the PSS and static groups, respectively. **(E)** Biosynthesis of MUFA depicts that SCD1 catalyzes the rate limiting step for the conversion of saturated FA (palmitic and stearic) to monounsaturated FA (palmitoleic and oleic). **(F)** Compound correlation network was generated with Cytoscape software. SCD1-mediated metabolites are interconnected with the glycolytic metabolites. Organic layout was used to organize the network according to major functional classes. The cluster predominantly involved with metabolites and showed significantly changed and are indicated in green nodes (palmitate, oleic acid and azelaic acid). A highly connected network within the metabolites cluster reveals the effect of exercise on mice highlighting the interactions with desaturase metabolism and other pathways.

### Exercise-augmented PSS mitigated flow recirculation in the aortic arch

To obtain the boundary conditions for our *in-silico* analysis of exercise-augmented PSS, we interrogated the mouse aortic arch via an ultrasound transducer (**Figure 2A**). Electrocardiogram (ECG)-gated Pulsed wave (PW) Doppler was acquired for the time-dependent inlet flow boundary condition (**Figure 2C**). This inlet boundary condition was calibrated for the exercise model as previously reported ^29, 30^. We demonstrated the B-mode image of the aortic arch, and used three-element wind Kessel model for the 4 outlet boundary conditions; specifically, the brachiocephalic, the left common carotid, and the left subclavian artery ^31^ (**Figure 2B**).

**Figure 2.**
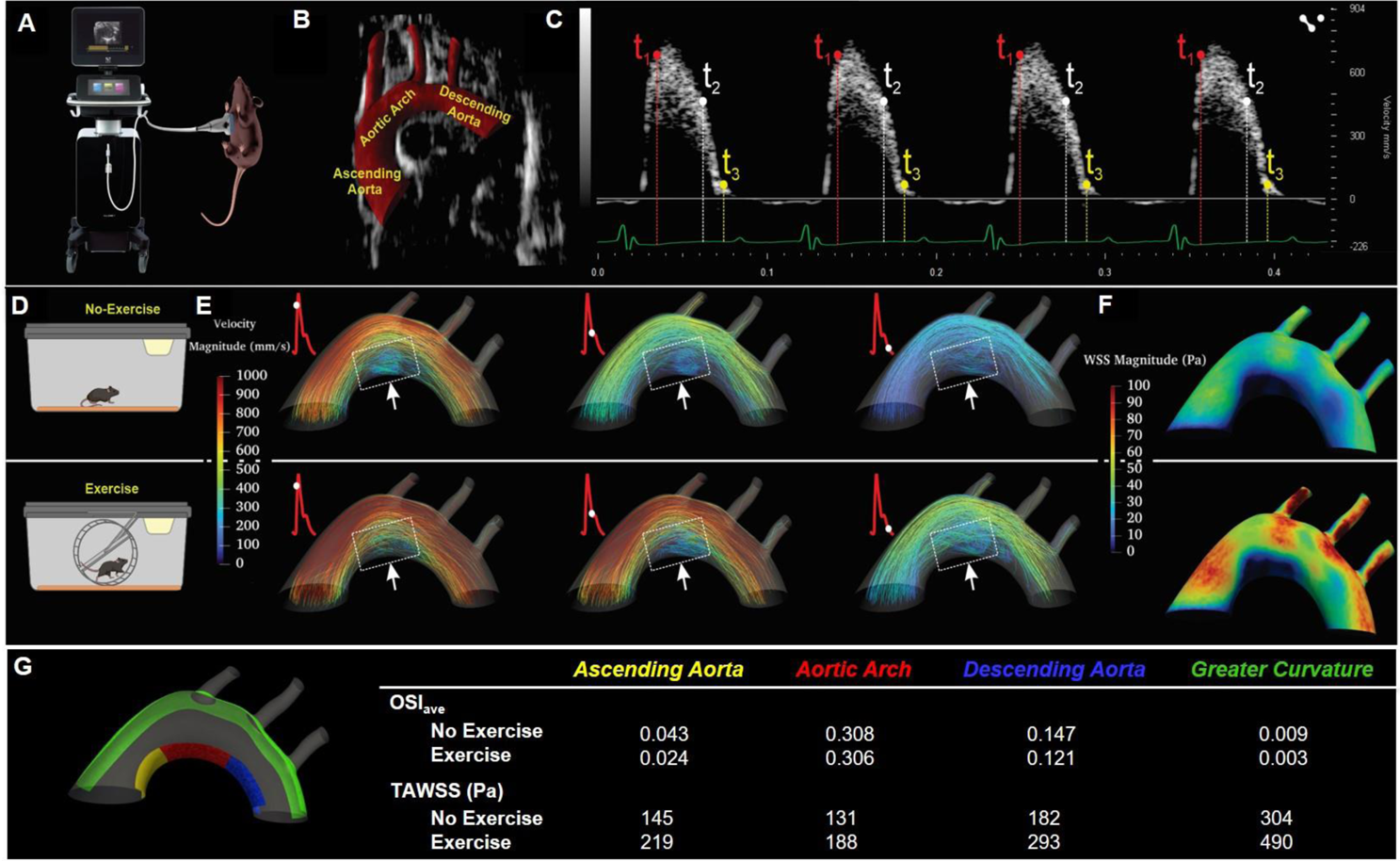
*In silico* analysis of exercise-augmented PSS. 4-D CFD simulation was performed to analyze hemodynamic profiles in the mouse aorta in response to exercise. **(A)** A schematic of ultrasound B-mode and PW Doppler imaging were acquired from the anesthetized mouse. **(B)** A representative B-mode image of the mouse aortic arch and the three branches, and the descending aorta. **(C)** PW Doppler measurement was acquired from the mouse ascending aorta over 4 ECG-gated cardiac cycles. **(D)** Schematic of mouse engaging in voluntary wheel running. **(E)** Velocity-colored streamlines along the aortic arch demonstrating disturbed flow developed at the lesser curvature, whereas exercise-augmented shear stress mitigates flow recirculation at the disturbed flow-prone lesser curvature. **(F)** TAWSS over several cardiac cycles was computed and compared between no exercise and voluntary wheel running. Note that pulsatile shear stress develops at the greater curvature of the aortic arch and descending aorta, whereas disturbed flow, including oscillatory shear stress, develops at the lesser curvature. Exercise-augmented pulsatile shear stress mitigates disturbed flow at the lesser curvature. **(G)** Comparison between OSI and TAWSS across color-coded regions of the lesser and greater aortic curvature, with or without exercise.

Four-D (space + time) Computational Fluid Dynamics (CFD) revealed the velocity-colored streamlines over a representative cardiac cycle (**Figure 2E**). Our time-dependent CFD simulation demonstrated the peak of the velocity magnitude profile proximal to the greater curvature, and low WSS at the lesser curvature. Furthermore, our model captured the flow recirculation zone developed at the lesser curvature (dotted square boxes) during systole (**Figure 2E**). We further simulated the exercise-augmented blood flow by applying the heart rate and flow rate variation to the model. Our results showed mitigation of the flow recirculation zone at the lesser curvature of the aortic arch. The comparison between sedentary (No EX) and voluntary wheel running conditions indicated an upward trend for the time-averaged wall shear stress (TAWSS or *τ*_ave_) at different regions of the aortic arch. The time-averaged oscillatory shear index (OSI_ave_) was calculated and compared for the sedentary and exercise groups (**Figure 2G** and **Supplemental Figure 5)**. The OSI_ave_ on the surface indicated that oscillatory flow along the lesser curvature of the aortic arch was mitigated in response to exercise. While it remains experimentally challenging to acquire real-time hemodynamic profiles, our *in-silico* analysis supports that exercise modulates PSS (*τ*_ave_ and OSI_ave_) to mitigate flow recirculation in the disturbed flow-prone lesser curvature.

### Exercise activated endothelial SCD1 in the ER to catalyze lipid metabolites

Wild-type C57BL/6J mice were subjected to voluntary wheel running to recapitulate exercise-augmented PSS (**Figure 3A and 3B**). Plasma samples were collected at baseline and after 24 h of exercise. Untargeted metabolomic analysis revealed that OA was markedly increased, while PA exhibited an increasing trend although it did not reach the level of statistical significance (**Figure 3C and 3E**). Saturated palmitic and stearic acids were unchanged (**Figure 3D**). PCA revealed a distinct separation of lipid metabolites after exercise (**Figure 2F**). Consistent with the finding in PSS-induced HAECs, PSS increased glycolytic metabolites including lactate, fumarate, oxalate, and succinate, as previously reported ^25^ (**Supplemental Figure 1A**).

**Figure 3.**
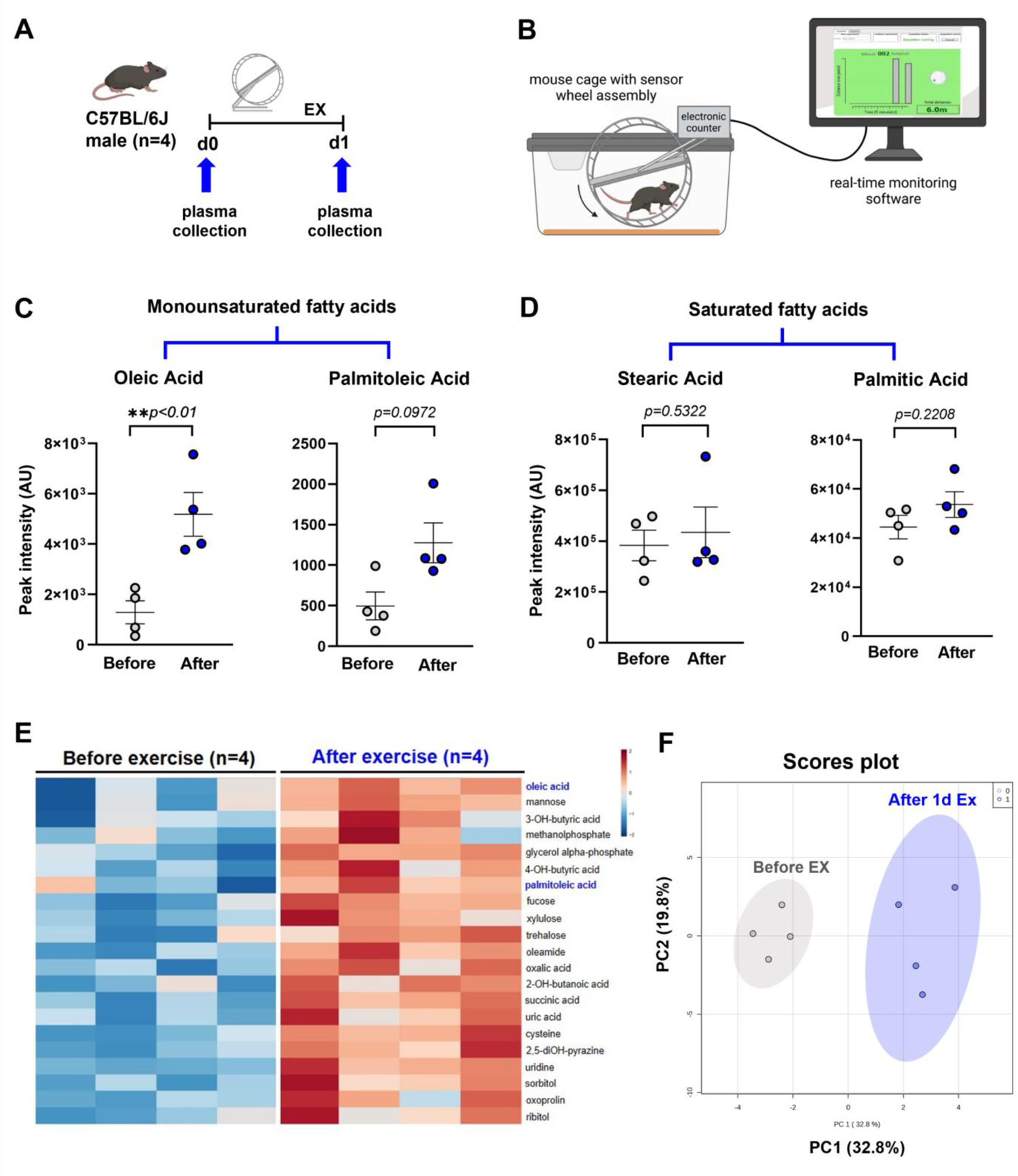
Exercise activates SCD1 to catalyze lipid metabolites. **(A)** A schematic representation of the animal exercise protocol. Plasma was collected from male wild-type C57BL/6J before and after 24 h of voluntary wheel running for metabolomic analysis (n=4 mice for each group). **(B)** The voluntary wheel running system provided real-time monitoring of rodent running activity (figure generated with BioRender). (**C)** 24 hours of exercise increased oleic, palmitoleic and linoleic acids; however, the latter two metabolites did not reach statistical significance (oleic acid: \*\**p* < 0.01 vs. before exercise, n=4). **(D)** Exercise did not change the plasma levels of saturated FA stearic and palmitic acid. **(E)** Heatmap of plasma metabolites before and after exercise reveals that OA was elevated in all 4 mice after normalization and scaling using the Pareto method. **(F)** Score plots by the principal component analysis support the 90% confidence intervals before and after exercise.

We next performed *in situ* hybridization (RNAscope) with a *Scd1-*specific probe, revealing prominent *Scd1* mRNA expression in the endothelial layer of both the aortic arch and descending aorta after 2 weeks of exercise (**Figures 4, A-C)**. *En face* immunostaining of the exposed aortic endothelium demonstrated an increase in the perinuclear expression of SCD1 but was absent in *Scd1*^EC-/-^ mice (**Figure 4D and 4E**), supporting the subcellular localization of SCD1 in the ER. Changes in body weight and exercise parameters were quantified for both female and male C57BL/6J mice (**Supplemental Figure 1B** and **Supplemental Table 1)**. Taken together, our findings support the interpretation that exercise-augmented PSS up-regulates SCD1 in the endothelial ER to catalyze lipid metabolites. Furthermore, our *in vitro* and *in vivo* findings support the previous report that PSS-activated *Scd1* is a *Pparγ* target gene ^23^ (see **Supplemental Figure 2, A-E**).

**Figure 4.**
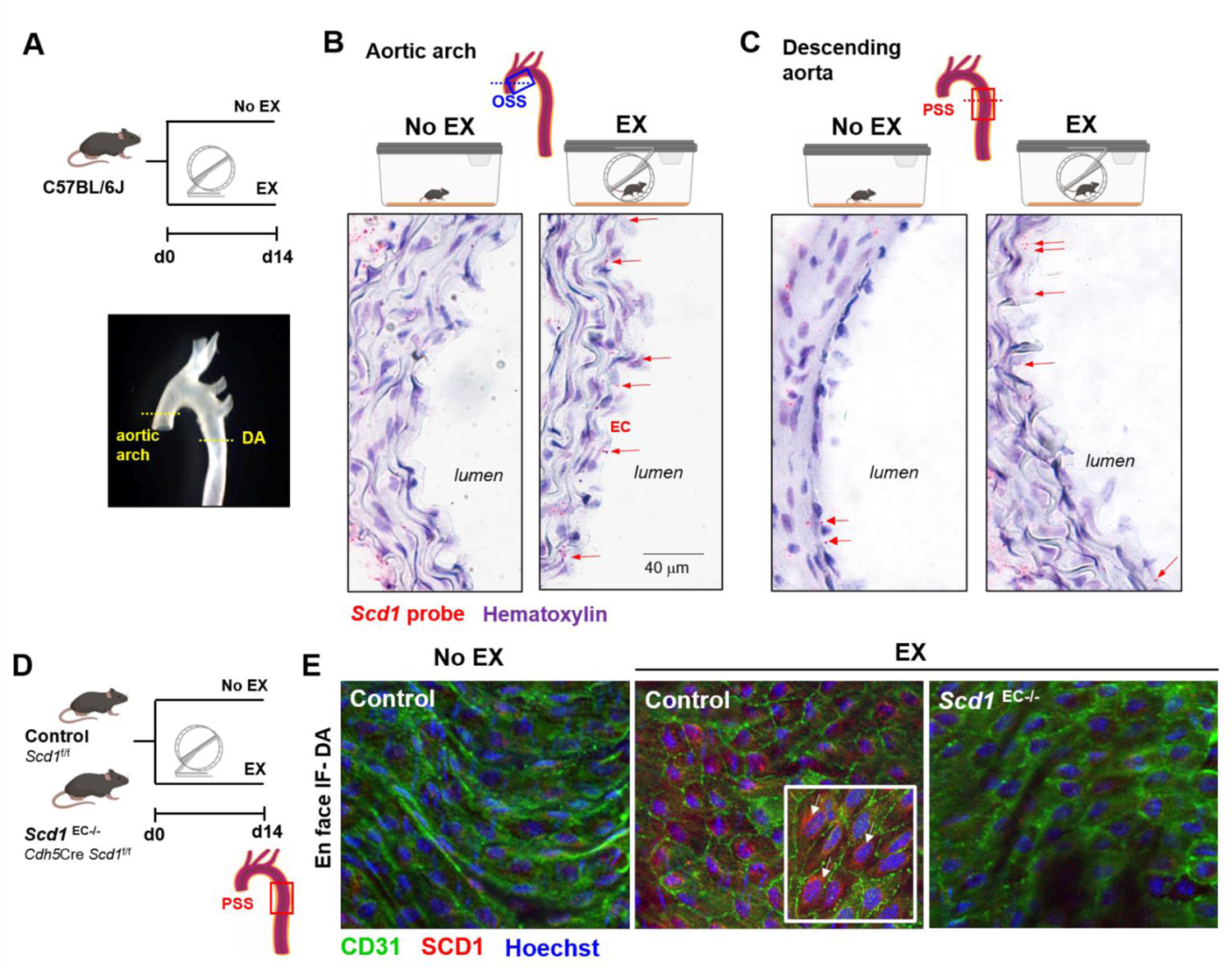
Exercise activates endothelial SCD1 in the endoplasmic reticulum (ER). **(A)** Wild type C57/BL6J mice underwent 14-day voluntary wheel running protocol. The aortas were processed for transversal cryosections at the level of the aortic arch and the descending aorta (DA). **(B and C)** The *Scd1*-specific probe (RNAscope) demonstrates prominent endothelial staining following 14 days of exercise-induced PSS both in the flow-disturbed aortic arch (B) and the DA (C). Red dots (arrows) indicate the mRNA molecules in the intima layer. *Scd1* was also present in some smooth muscle cells and periaortic adventitia (adv) in both groups. Microphotographs were taken at 63X magnification. **(D)** The endothelial-specific *Scd1* deleted mice underwent 14-day exercise protocol. **(E)** *En face* immunostaining of exposed aortic endothelium reveals that exercise increased the SCD1 expression (red) in the aortic endothelial cells (CD31+, green) (n=3). A higher magnification image (63X: right lower corner insert) showed perinuclear SCD1 staining (arrows), supporting the subcellular localization of SCD1 in the ER. No staining was observed in the *Scd1* ^EC-/-^ mice undergoing exercise.

### Endothelial-specific SCD1 deletion abrogates the SCD1-mitigated anti-inflammation and atherosclerosis in the disturbed flow-prone aortic arch

*Ldlr*^-/-^ mice with or without conditional deletion of the *Scd1* gene upon Cre-mediated recombination were generated under the endothelial-specific promoter Cdh5 (VE-cadherin) (*Ldlr*^-/-^*Scd1*^EC -/-^) ^32^. Mice received high-fat diet (HFD) for 28 days to promote the development of fatty streak in the aorta ^33^. After the first 14 days of HFD, some mice underwent voluntary wheel running for 14 days while continuing on HFD (HFD-EX) (**Figure 5A**). Body weight gain and exercise data are included in **Supplemental Figure 3** and **Supplemental Table 2**.

**Figure 5.**
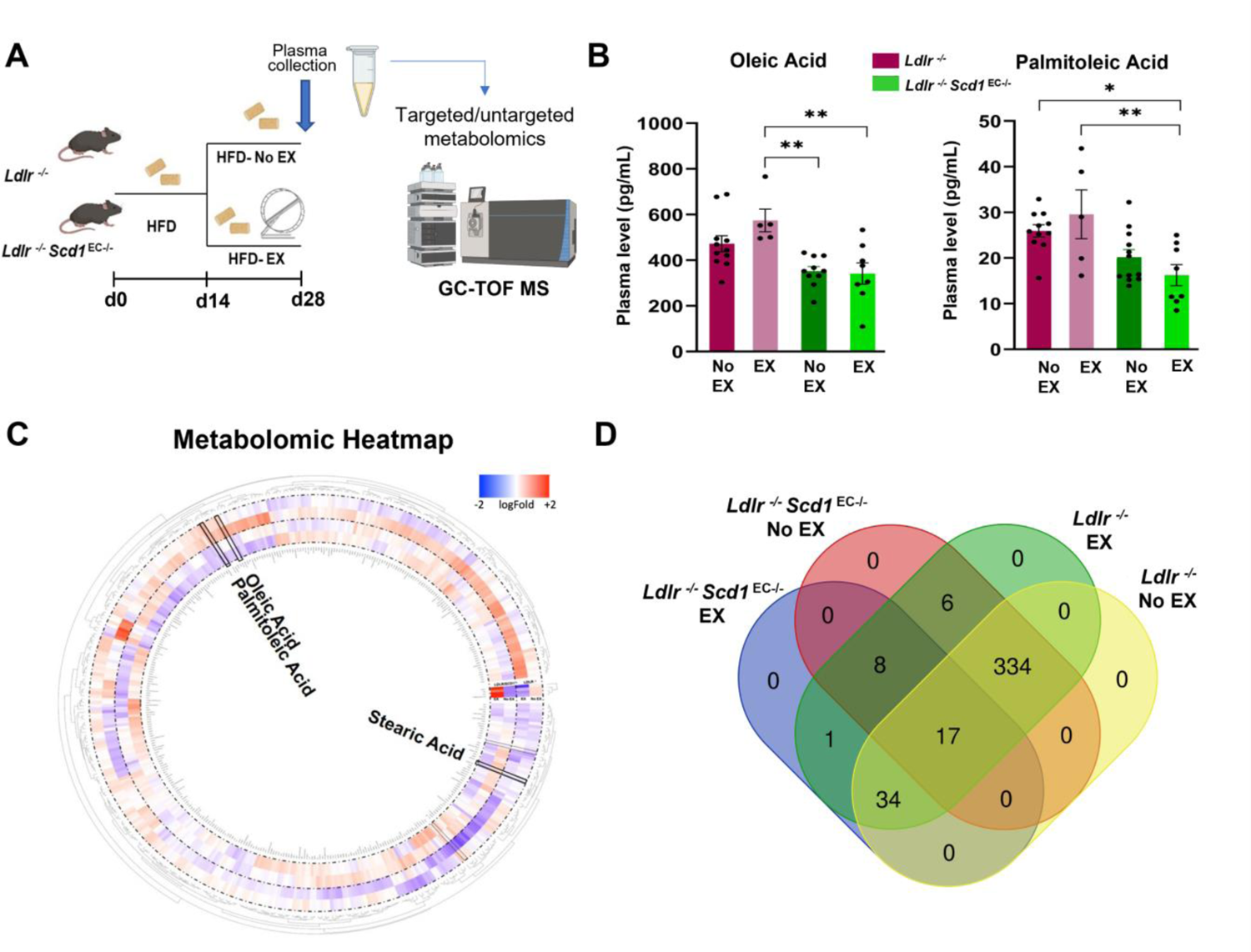
Exercise-mediated lipid metabolites, OA and PA, are reduced in mice with endothelial-specific SCD1 deletion. **(A)** The exercise protocol depicts the *Ldlr*^-/-^ mice that were crossed with *Cdh5Cre*; *Scd1*^flox/flox^ to generate endothelial-specific SCD1 deletion (*Ldlr^-/-^ Scd1*^EC-/-^). Adult mice at 8 weeks received HFD for 14 days. The mice were then divided into exercise (EX) and no exercise (No EX) groups for additional 14 days of exercise. **(B)** The absolute plasma concentration of OA was elevated after exercise in the *Ldlr*^-/-^ mice, whereas OA concentration was reduced and remained reduced after exercise in the *Ldlr^-/-^Scd1*^EC-/-^ mice. The plasma concentration of PA also followed the similar trends, suggesting that exercise activates SCD1 to catalyze OA and PA (\**p* < 0.05, *p* < 0.01; n=4-12; each dot represents an individual animal). **(C)** Circular heatmap of untargeted metabolomic analysis capturing known and unknown ID metabolites highlights changes in OA and PA further supporting that SCD1 deletion resulted in a decrease in both OA and PA, after normalization and scaling using the Pareto method. Circular heatmap was generated with Circos R package. **(D)** Venn diagram was generated with webtool Venn and it summarizes the shared metabolites among different conditions.

On day 28, quantitative and targeted metabolomic analyses revealed that the basal levels of OA and PA in plasma were reduced in the *Ldlr*^-/-^*Scd1*^EC -/-^mice as compared to*Ldlr*^-/-^ alone. After exercise, both OA and PA levels were significantly elevated in the *Ldlr*^-/-^ but not in the *Ldlr*^-/-^*Scd1*^EC-/-^mice (**Figures 5B and 5C, Supplemental Figure 4).** This finding confirms that endothelial-specific *Scd1* deletion abrogates exercise-activated lipid metabolites, including OA and PA. In addition, a subset of metabolites were differentially modulated under our experimental conditions (**Figure 5D**).

Ultrasound-based 4-D CFD simulation revealed disturbed flow along the inner curvature of the aortic arch, whereas pulsatile flow was prominent in the thoracic region of the descending aorta (**Figure 6A and Supplemental Figure 5)**. Confocal imaging of the aortic endothelium *en face* after immunostaining with anti-ERG (an endothelial nuclear marker) ^34^ and VCAM1 (an early atherosclerosis lesion marker) ^35^ revealed the presence of VCAM1 expression in the disturbed flow-prone inner curvature of the aortic arch in *Ldlr^-/-^* mice. *Ldlr*^-/-^ mice with endothelial-specific *Scd1* deletion (*Scd1*^EC-/-^) also displayed VCAM1 expression in the inner curvature. However, the 14-day exercise period mitigated levels of VCAM1 expression in *Ldlr*^-/-^ but not in *Ldlr*^-/-^ *Scd1*^EC-/-^ mice (**Figure 6B**). In the descending aorta, where the endothelium is exposed to PSS, deletion of endothelial *Scd1* led to an increase in VCAM1-positive endothelial cells compared to *Ldlr^-/-^* mice, and exercise did not mitigate the number of VCAM1 positive endothelial cells (**Figure 6C**)*. In situ* hybridization with a *Scd1*-specific probe corroborated prominent endothelial staining following exercise-induced PSS in the disturbed flow-prone aortic arch of *Ldlr*^-/-^ mice, which is absent in *Ldlr^-/-^Scd1*^EC-/-^ mice (**Figure 6E**). Taken together, exercise-augmented PSS is implicated in activating endothelial SCD1 for atheroprotection in the disturbed flow-prone vasculature.

**Figure 6.**
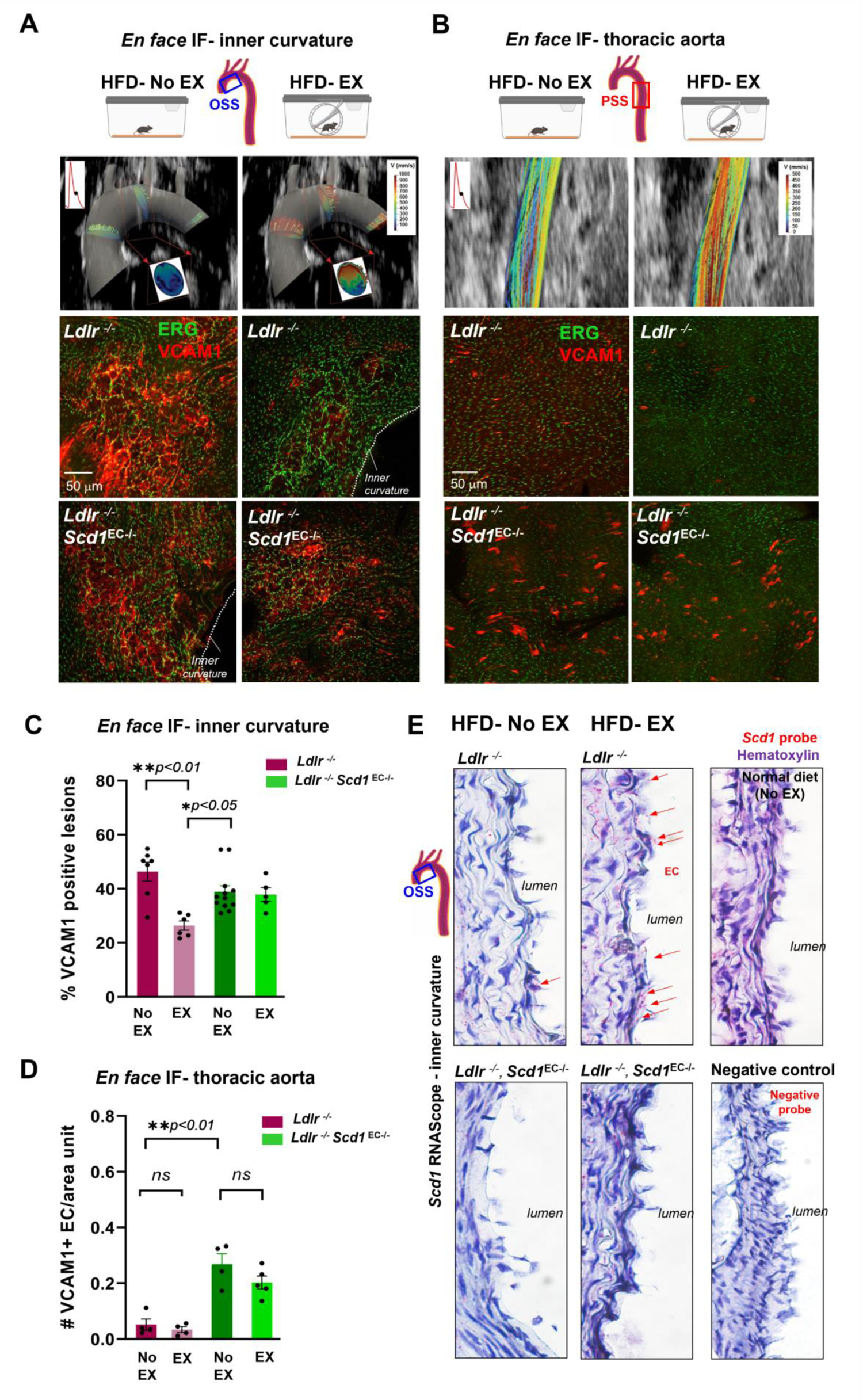
Endothelial-specific SCD1 deletion abrogates exercise-mediated anti-inflammatory mediators. **(A and B)** CFD simulates the spatial and temporal variations in wall shear stress in the aortic arch and descending aorta of a representative wild-type mouse. The ultrasound B-mode was used to reconstruct the 4D shear stress profiles. The cross-sections of the aortic arch illustrate the high shear stress localized to the greater curvature and low shear stress to the lesser curvature. Exercise-augmented shear stress mitigates disturbed flow at the lesser curvature. The CFD reconstruction over a cardiac a cardiac cycle (t_1_, t_2_, and t_3_) is included in Supplemental Figure 4; only t_2_ is depicted here. The colored streamlines indicate the CFD-derived velocity profiles along the aortic arch. The representative confocal Z-stack for the maximal projection of *en face* immunostaining with anti-ERG and anti-VCAM-1 in the inner curvature of the aortic arch and the thoracic region of the descending aorta. In the lesser curvature, the VCAM-1 staining was attenuated following exercise in the *Ldlr*^-/-^ mice but remained prominent in the *Ldlr^-/-^Scd1*EC-/- mice. In the thoracic aorta, the VCAM-1 staining was reduced with or without exercise in the *Ldlr*^-/-^ mice but were elevated with or without exercise in the *Ldlr^-/-^Scd1*^EC-/-^ mice. This finding further supports the interpretation that PSS activates endothelial SCD1 to mitigate arterial inflammation. **(C and D)** Quantification of *en fac*e immunostaining shown in **A** and **B** panels, respectively. Each dot corresponds to an individual animal (n=4-12); both the male and female data were pooled together. **(E)** The SCD1-specific probe (RNAscope) corroborates prominent endothelial staining following exercise-augmented PSS in the disturbed flow-prone aortic arch in *Ldlr*^-/-^ mice, which is absent in *Ldlr^-/-^Scd1*^EC-/-^ mice. *Scd1* expression remains unchanged after HFD as compared to normal chow diet. Red dots (arrows) indicate the mRNA molecules in the intima layer. Staining with a negative control probe shows complete absence of red positive dots. Microphotographs were taken at 63X magnification.

### SCD1-catalyzed oleic acid mitigates pro-inflammatory and ER stress mediators

We assessed the effects of OA on cultured HAECs in terms of *Vcam1*, *Icam1*, *Mcp1*, *Esel* (E-selectin) and the *Cxcl8* chemokine mRNA expression (**Figure 7A**). OA reduced these pro-inflammatory mediators in a dose-dependent manner ^35–37^. Also, OA decreased the expression of ER stress-related transcription factors, *Atf3*, *Atf4*, and *Atf6* **(Supplemental Figure 6).** These changes are consistent with our analysis of total liver gene expression after 14 days of exercise **(Supplemental Figure 7).** Furthermore, a recombinant adenovirus-based system was developed to overexpress SCD1 in HAECs via the *Cdh5* (CD144) promoter, which was validated by immunofluorescence (IF) and Western Blot. SCD1 overexpression mitigated inflammatory mediators in HAECs, such as *Cxcl8* and its receptor *Cxcr1* and the olfactory receptor 2 (*Or6a2)* ^38, 39^ (**Figure 7, B-D)**.

**Figure 7.**
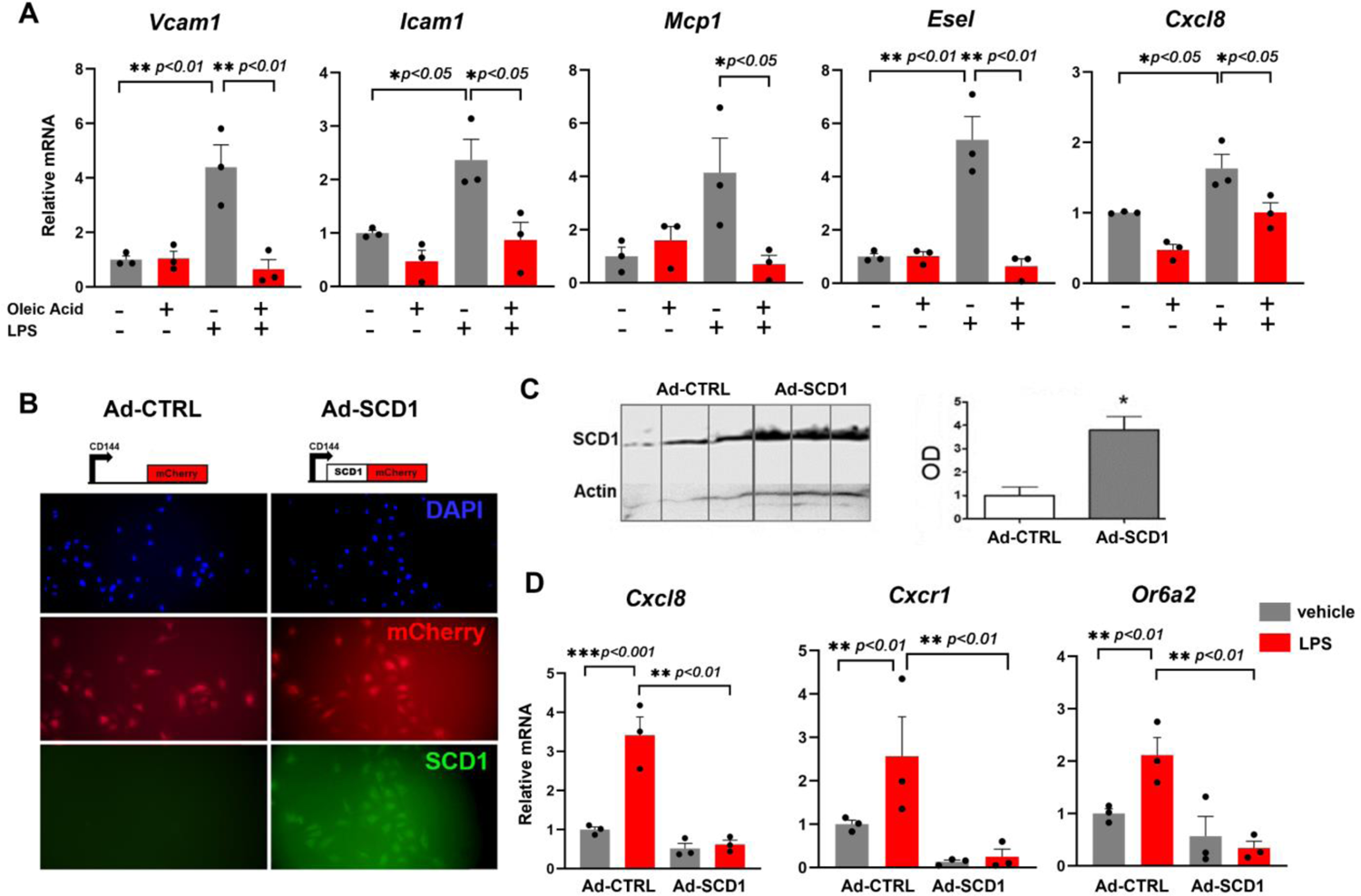
Oleic acid treatment or SCD1 overexpression mitigate pro-inflammatory mediators in HAECs. **(A)** HAECs monolayers were treated for 4 hours with OA at 0.2 mM in the absence or presence of LPS (20 ng/mL) and gene expression levels were analyzed by qRT-PCR. **(B)** HAECs were infected with adenovirus control (left panels) or SCD1 (right panels) and imaged for mCherry reporter and SCD1 expression. (C) Western Blot was performed and quantified. **(D)** Ad-SCD1 mitigated the LPS (20 ng/mL)-induced expression (4 hours) of *Cxcl8, Cxcr1* and *Or6a2* (n=3, \**p*<0.05, \*\**p*<0.01, \*\*\**p*<0.001).

### *Scd1* and gene interaction networks in response to biomechanical forces through single cell transcriptomic analysis of the mouse aorta

To further assess transcriptomic profiles of exercise-mediated endothelial gene expression in the aorta, we performed single-cell RNA sequencing on aortas from *Ldlr^-/-^* mice subjected to the following treatments for 4 weeks: 1) normal chow diet; 2) HFD; 3) HFD + EX (**Figure 8A**). Unbiased clustering analysis of transcriptional profiles identified 7 lineages based on cell-type specific markers (**Figure 8B and 8C, Supplemental Figure 8)**. 3D Volcano Plot further established *Scd1* as a gene differentially modulated by exercise in the aortic endothelium (**Figure 8D**). As a corollary, by integrating our endothelial transcriptomic data with a published database ^40^ using Cytoscape we uncovered potential candidate genes interacting with *Scd1* in ECs, including adiponectin receptor 2 (*Adipor2*) ^41^, peroxisomal acyl-coenzyme A oxidase 1 (*Acox1*) ^42, 43^ and insulin receptor substrate 2 (*Irs2*) ^44–46^ (**Figure 8E**). Furthermore, gene-metabolite interconnection networks revealed a relationship between *Scd1* and the *Acsl* family of long-chain acyl CoA synthetase genes and acyl-CoA thioesterase genes (*Acot*), which are involved in intracellular lipid metabolism.

**Figure 8.**
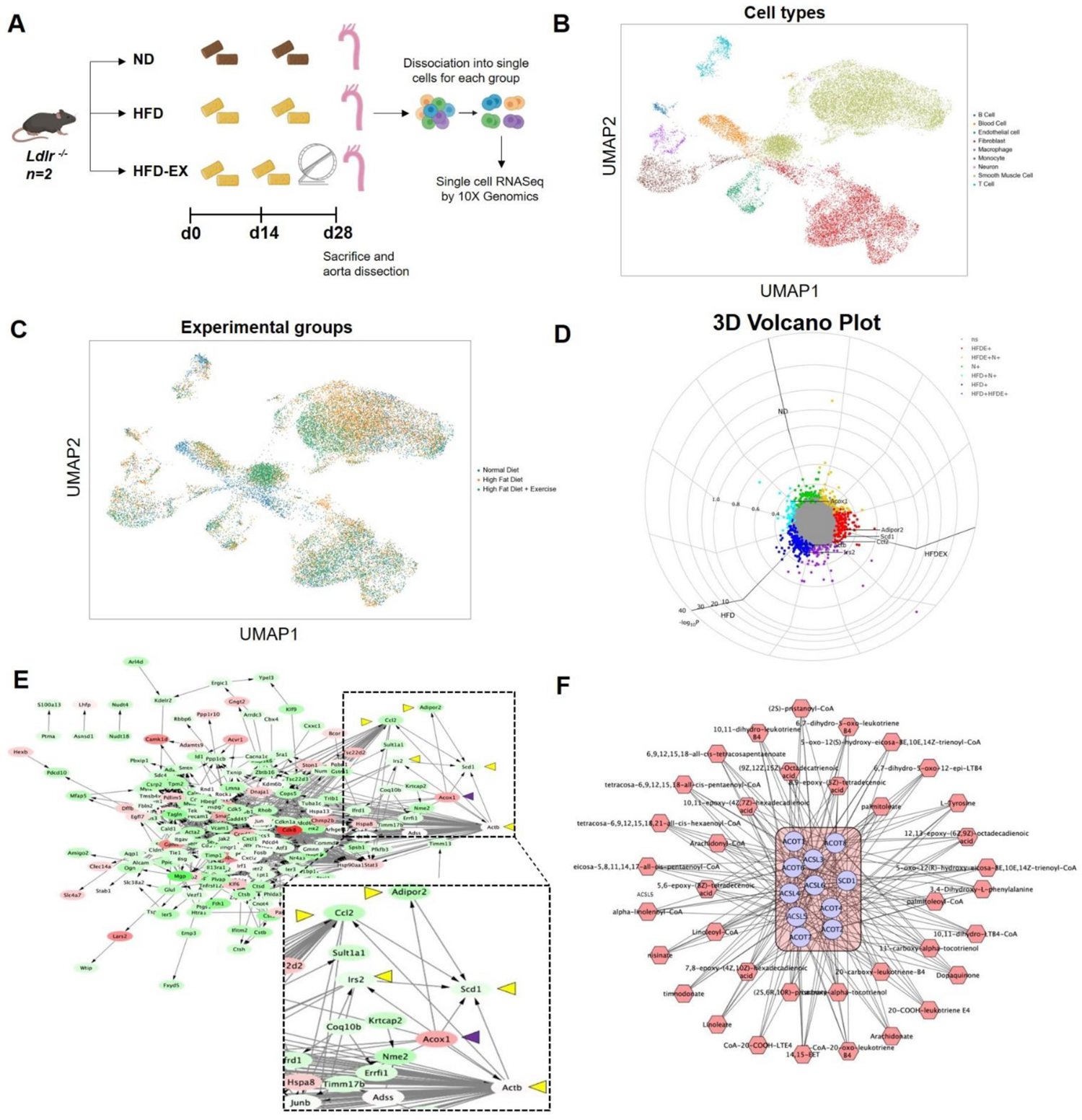
Single cell transcriptomics of mouse aorta and network interactions. **(A)** Schematic diagram of the experimental protocol. *Ldlr*^-/-^ mice (n=2 per group) were subjected to normal chow diet (ND, Control), HFD or HFD plus exercise (HFD-EX) during the last 2 weeks of the treatment. Aortas were collected after 28 days. A total of 20,000 cells were analyzed. **(B and C)** Representative UMAP of cellular clusters in the mouse aorta. **(D)** 3D cylindrical volcano plot of differentially expressed genes comparing ND vs HFD vs HFD-EX was created by using Volcano 3D R package. Vectors for sample mean Z score per gene were projected onto a polar coordinate space analogous to RGB (red-green-blue) color space mapping to HSV (hue-saturation-value) as described in ^94^. **(E)** Gene network generated with Cytoscape software from EC transcriptomics and published database of mechanosensitive genes ^40^. **(F)** Analysis of interaction networks between endothelial transcriptomic and plasma metabolomics data highlighting the connection of *Scd1*, *Acsl* and *Acot* gene families with several exercise-induced metabolites.

## DISCUSSION

Vascular diseases, including coronary artery disease, stroke, and peripheral arterial disease, predispose individuals to develop chronic disability and increase the healthcare burden ^47^. Exercise intervention is an effective lifestyle modification to ameliorate cardiometabolic disease, however, the underlying flow-sensitive metabolomic transducers that act to ameliorate disease are not completely known. In this study, we demonstrate how exercise activates SCD1 to catalyze vascular protective lipid metabolites in the disturbed flow-prone region of the aortic arch. By integrating our dynamic flow system, metabolomic analyses, ultrasound imaging to reconstruct computational domain for CFD, and wheel running system, we demonstrated that exercise augments PSS (*τ*_ave_ and OSI_ave_) to activate endothelial SCD1 in the disturbed flow-prone aortic arch (**Figure 9**). Furthermore, endothelial-specific SCD1 deletion or overexpression established that exercise-activated SCD1 catalyzes the lipid metabolite OA, to attenuate inflammatory and atherosclerotic mediators in the lesser curvature of the aortic arch.

**Figure 9.**
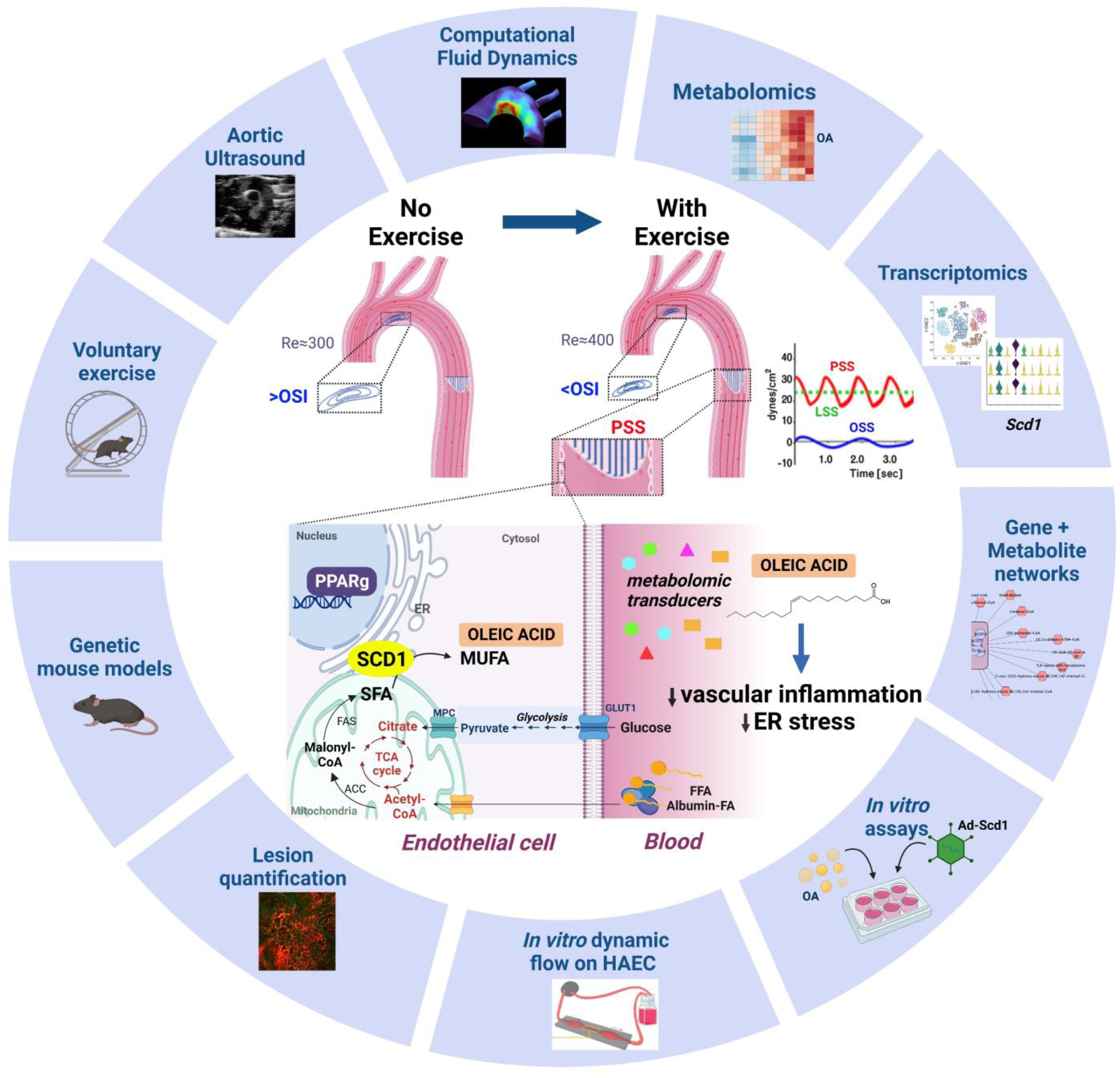
Working model for exercise-induced PSS in the aorta and activation of the flow-responsive PPARγ-SCD1 pathway for aortic atheroprotection. Exercise-induced PSS reduces flow recirculation and OSI in the aortic arch. At the cellular level, shear stress-mediated glycolytic flux yields pyruvate as a substrate for FA metabolism. EC can uptake and metabolize either free fatty acids (FFA) or albumin-bound FAs. SCD1 located in the ER membrane adjacent to the mitochondrial membrane is the limiting step in the conversion of SFA into MUFA. The SCD1 product oleic acid reduces NF-κB-induced inflammation and ER stress genes. MPC: mitochondrial pyruvate carrier; TCA: tricarboxylic acid. ACC: acetyl-CoA carboxylase; FAS: fatty acid synthase; Re; Reynolds number. Figure was generated with BioRender.

Endothelial cells line the lumen of the vasculature and regulate the transport of oxygen and nutrients. Hemodynamic shear stress modulates cell metabolism to maintain endothelial homeostasis ^48, 49^, and PSS is well-recognized to be atheroprotective, by attenuating pro-inflammatory mediators and NADPH oxidase system ^10, 12, 50, 51^. To identify PSS-mediated endothelial metabolites, we seeded the HAECs in our custom-designed in vitro flow system to mimic the spatial and temporal variations in WSS. Our metabolomic analysis showed that PSS increased SCD1-catalyzed fatty acid metabolites in HAECs, and this observation guided us to investigate exercise-activated SCD1 as a metabolomic transducer to mitigate inflammatory mediators and ER stress.

SCD1 is an enzyme anchored in the ER that catalyzes the conversion of saturated long-chain fatty acids, palmitic (16:0) and stearic (18:0), to MUFAs palmitoleic (16:1) and oleic (18:1). The resulting MUFAs are the major components of triglycerides, cholesterol esters, and phospholipids ^24, 52^. Using subcellular fractioning, previous studies indicated that SCD1 is localized in the mitochondria-associated membrane (MAM), a specific region of the ER in close proximity to the outer mitochondrial membrane that is enriched in enzymes responsible for phospholipid and triglyceride biosynthesis ^53^. SCD1 is expressed in a broad range of tissues and at high levels in the insulin-sensitive hepatocytes and white and brown adipocytes ^54, 55^. Other tissue-specific SCD isoforms have also been identified, including SCD2, which is found mainly in the brain ^56^, SCD3 in the skin and sebaceous gland ^57^ and SCD4 in the heart ^58^. However, endothelial SCD1 has not been explored *in vivo*, primarily because of experimental challenges and low basal expression compared to highly metabolically active tissues.

The expression of SCD isoforms is highly regulated by multiple factors, including diets and hormones ^59^. SCD1 is expressed in multiple organs and tissues to catalyze lipid metabolites in association with adipose, cancer cell, and skeletal muscle metabolism, as well as liver and lung fibrosis ^60–62^. Global deficiency of SCD1 results in reduced adiposity, increased insulin sensitivity, and resistance to diet-induced obesity ^63^. Interestingly, the global deletion of SCD1 in *Ldlr* null mice leads to abnormal lipid metabolism, ER stress, and atherosclerosis ^64^. Furthermore, hand grip strength, duration of swimming time to exhaustion, and voluntary vs. forced treadmill wheel running support the notion that SCD1 overexpression increases cardiovascular endurance in mouse models ^65^.

In this context, we developed the endothelial-specific *Scd1* deletion in *Ldlr* null mice. During exercise, autonomic function elevates both blood pressure (BP) and heart rate (HR); thus, leading to increased blood flow and shear stress ^66^. By deploying micro shear stress sensors into the rabbit aorta, we demonstrated a positive correlation between BP/HR and pulsatile shear stress ^67^. Accordingly, wild-type mice undergoing 2 weeks of wheel running developed PSS-activated endothelial SCD1 expression, as evidenced by in situ hybridization and *en face* immunostaining of the aortas. This data indicates that exercise-augmented PSS activates endothelial SCD1 expression.

The cardiovascular benefits of exercise have been well-documented to improve endothelial function and ameliorate both vascular inflammation and oxidative stress ^68^. Exercise elevates the plasma levels of serum clusterin that increases plasticity and reduces inflammation in the cerebrovascular system ^8^. Recently, exercise was reported to increase both the human and racehorse plasma metabolite, *N*-lactoyl-phenylalanine (Lac-Phe), which also suppressed feeding and obesity in mice ^69^.

Despite these salutary effects, regular exercise is limited to certain populations due to physical disabilities and/or socioeconomic limitations ^70^.The prevalence of sedentary lifestyles remains high in industrialized nations, including the United States. To tackle this problem, the National Institutes of Health launched the Molecular Transducers of Physical Activity Consortium (MoTrPAC) with the aim to integrate large-scale multi-omic analyses with bioinformatics to identify molecular transducers as pharmacologic interventions to mimic or enhance the therapeutic effects of exercise ^16^. Several signaling pathways have been identified to mediate exercise’s benefits, including IGF1/PI3K/AKT, C/EBPβ/CITED4, PPAR family, PGC1α, AMPK, eNOS/NO, and certain noncoding RNAs ^71^.

PPARs belong to the nuclear receptor superfamily of ligand-activated transcription factors regulating lipid and glucose metabolism, energy homeostasis and inflammation ^72^. Previous studies have demonstrated that shear stress induces PPARα, δ and γ to regulate endothelial function ^73, 74^. PPARγ agonists belong to the thiazolidinedione class of drugs such as troglitazone and rosiglitazone that attenuate palmitate-induced ER stress and apoptosis by activating SCD1 induction in macrophages ^75^. PPARγ was shown to protect against IL-1β-mediated endothelial dysfunction through a reduction of oxidative stress responses ^76^ and is essential for preventing aging-associated endothelial dysfunction ^77^. Our flow system revealed that PSS-activated SCD1 expression in HAEC was PPARγ-dependent, consistent with the previous report that laminar shear stress up-regulated SCD1 through a PPARγ-specific mechanism ^23^. To further demonstrate the PPARγ-dependent endothelial SCD1 expression *in vivo*, we subjected transgenic mice expressing the dominant-negative V290M PPARγ mutation (E-V290M Tg) ^78^ to voluntary wheel running. Immunostaining for SCD1 was absent in the aortic endothelial cells as compared to the wild-type **(Supplemental Figure 2E),** corroborating the PPARγ-dependent SCD1 activation.

*Ldlr*^-/-^ mice fed with HFD develop vascular lesions in a time-dependent manner and have been widely used to mimic human atherosclerosis. The atherosclerotic lesions develop at susceptible regions associated with low shear stress or disturbed flow, such as the inner curvature and branching points of the aorta ^79^. Hemodynamic-driven disruptions of the integrity of the arterial intima drive atherogenic inflammation, associated with increased VCAM1 expression, recruitment of Ly6G-positive neutrophils and subsequent accumulation of erythrocyte-derived iron and lipid droplets, which form the incipient atherosclerotic lesions known as fatty streaks ^35^. In our study, VCAM1 expression was prominent in the inner curvature of the aorta in *Ldlr*^-/-^ mice after 4 weeks of HFD but absent in the PSS-mediated descending aorta. However, *Ldlr*^-/-^ mice with endothelial-specific SCD1 deletion developed more pronounced VCAM1 positive areas in the descending aorta. After 14 days of exercise, the VCAM1 positive lesions were attenuated in the *Ldlr*^-/-^ but not *Ldlr*^-/-^*Scd1*^EC-/-^ mice, further supporting the notion that exercise-activated SCD1 mitigates inflammatory mediators.

Diets enriched in saturated fatty acids are linked to increased cardiovascular disease risk, whereas monounsaturated fatty acids have been associated with improved cardiovascular outcomes ^80^. OA is the predominant MUFA in the Mediterranean diets, including olive oil as the principal source of fat. There is strong evidence for an association between a Mediterranean-style diets and cardiovascular disease protection ^81^. Our *in vitro* data with OA treatment in HAECs support previous reports that OA inhibits liposaccharide (LPS)-induced endothelial oxidation ^82–84^, reduces NF-κB activation ^85^, and ameliorates insulin resistance ^83^. In our study, both OA treatment or SCD1 overexpression in HAECs reduced *Cxcl8* (interleukin-8), a member of the chemokine family with proinflammatory properties through CXCR1 and CXCR2 receptors ^38^. In addition, OA reduced the ER stress-related transcription factors. In this context, SCD1 deficiency predisposes the cells to ER stress which activates the unfolded protein response (UPR) by increasing ATF6 production, ER stress-mediated pathways, including BiP, CHOP, HSP90B1, and XBP1 ^86–88^. Endothelial SCD1 overexpression further reduced the *Or6a2* expression, for which the murine ortholog *Olfr2* is expressed in the mouse macrophages to promote atherosclerosis ^39^. Despite the beneficial effects of SCD1 overexpression *in vitro* in HAECs, a major hurdle for *in vivo* application of this system is sequestration of adenoviruses in the liver sinusoids ^89^.

As a corollary, our network analysis of endothelial transcriptomic profiles in response to HFD and exercise-induced biomechanical forces uncovered three candidate genes that interact with *Scd1* activity and deserve further mechanistic investigation. *Adipor2* regulates membrane fluidity by maintaining MUFA levels in HEK293 cells through sustained desaturase gene expression ^90^. *Acox1* protects against LPS-driven inflammation ^42^ and *Irs2* regulates eNOS phosphorylation and glucose uptake by skeletal muscle ^44–46^.

## Conclusion

Exercise-modulated hemodynamic shear stress (PSS: *τ*_ave_ and OSI_ave_) activates ER transmembrane SCD1 to catalyze vascular protective lipid metabolites in the disturbed flow-prone aortic arch. While SCD1 is associated with lipid metabolism across multiple tissues, we reveal that endothelial-specific SCD1 ameliorates inflammatory mediators and atherosclerosis in the lesser curvature of the aortic arch.

Overall, our comprehensive integration of genetic mouse models, exercise, CFD, light-sheet plaque assessment and bioinformatic network analysis of genes and metabolites provides new insights into the flow-sensitive endothelial transducer SCD1 and identifies lipid metabolites as therapeutic targets to mitigate cardiometabolic syndromes.

## METHODS

### Mice Lines

Male and female C57BL/6J mice from Jackson (Strain 000664) were used for the exercise studies. The endothelial dominant-negative *Pparγ* (EC-DN-*Pparγ*) mice line was kindly provided by Curt D. Sigmund at the University of Iowa ^78^. These mice express a human dominant negative *Pparγ* variant with the human V290 mutation under the control of the endothelial-specific *Cdh5* promoter.

Mice carrying the *Ldlr^tm1Her^* mutation were obtained from Jackson Laboratories (Strain 002207) to generate *Ldlr^tm1Her/tm1Her^* (*Ldlr* ^-/-^) mice. To study SCD1-mediated arterial inflammation, we crossed *Ldlr* ^-/-^ with mice carrying a conditional mutation of the *Scd1* gene (*Scd1^flox/flox^*) generated by James Ntambi lab at the University of Wisconsin Madison ^54, 91^. The endothelial deletion of *Scd1* gene was achieved by Cre recombination with *Cdh5Cre+* mice, in which the *Cdh5* promoter (also known as *VE-Cadherin*) drives the constitutive expression of *Cre* in endothelial cells. *Cdh5Cre* mice were obtained from the laboratory of Luisa Iruela-Arispe^32^. *Ldlr*^-/-^, *Scd1^flox/flox^*, *Cdh5^Cre^* mice (subsequently called *Ldlr*^-/-^, *Scd1*^EC-/-^) were generated through appropriate breeding schemes and PCR genotyping in tail DNA. Littermates *Ldlr*^-/-^, *Scd1^flox/flox^* (or *Ldlr*^-/-^ for simplicity) were used as controls. All mice were maintained on a C57BL/6J genetic background to reduce genetic variability.

### Induction of Atherosclerosis

Mice at 6-8 weeks of age were used for all our studies. Mice were fed a high fat diet (HFD, Teklad TD.88137, Envigo, Indianapolis, IN) for 4 weeks. The formulation is highly enriched in saturated fatty acids (>60% of total fatty acids) and contains 42% kilocalories from fat. The diet was supplied as soft pellets and replaced every 2-3 days to ensure freshness. After 2 weeks, some animals were transferred to individual exercise cages and continued receiving HFD for 2 more weeks (**Figure 5A**).

### Voluntary Wheel Exercise Studies

The wild-type C57BL/6J mice fed a regular rodent chow versus *Ldlr*^-/-^ and *Ldlr*^-/-^ *Scd1*^EC-/-^ mice on HFD were assigned to exercise. They were transferred to individual polycarbonate cages with a stainless-steel running wheel. Each wheel had an automated counter to record the distance and speed at which each animal ran in either direction (BIO-ACTIVW-SOFT from Bioseb, Pinellas Park, FL, USA). Spontaneous wheel running was monitored continuously over 2 weeks. Animals received food and water *ad libitum* during the data collection period. Mice in the no exercise group were housed in regular static cages in the same testing room to minimize environmental variations.

### Quantification of Arterial Strain and Wall Shear Stress after Exercise

We used the B-mode images of the aortic arch to simulate the time-dependent 3-D model, and reconstructed a 3-D model of the lumen structure by resolving the vessel wall curvature along the centerline of the aorta and its branches using our customized script (Matlab). To obtain the boundary conditions for our in-silico analysis of exercise-augmented PSS, we interrogated the mouse aortic arch via an ultrasound transducer (Vevo 3100, FUJIFILM VisualSonics) (**Figure 2A and 2C**). ECG-gated PW Doppler was acquired for the inlet boundary condition of the time-dependent flow of the aortic arch. This inlet condition was calibrated for the exercise model as previously reported ^29, 30^. The three-element Windkessel model was implemented at the 4 outlets of the model computational domain; namely, brachiocephalic, left common carotid, left subclavian artery, and descending aorta ^31^.

Time-averaged Wall shear stress 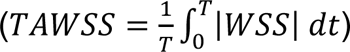 contours, oscillatory shear index 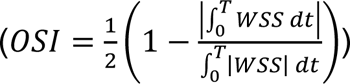 and velocity profiles for a time-dependent model were acquired by solving the Navier-Stokes equations using the using SimVascular’s svFSI solver ^92^.

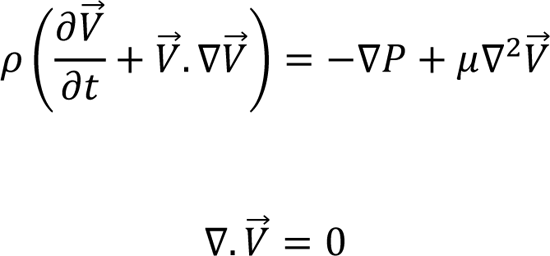

### A Dynamic Flow System to Simulate PSS Profiles

A dynamic flow channel was used to recapitulate hemodynamics of human carotid arterial bifurcations ^93^. The flow system was designed to generate well-defined flow profiles across the width of the parallel flow chamber at various temporal gradients (∂τ/∂t), frequencies, and amplitudes. Metabolite samples were collected from HAEC monolayers exposed to pulsatile shear stress (PSS, *τ*_ave_ = 50 dyne·cm^−2^ accompanied by ∂τ/∂t = 71 dyne·cm^−2^·s^−1^ at 1 Hz) or static conditions for 4 h. Cells were trypsinized, fixed in 4% PFA, and immediately stored at −80°C before submission for metabolomic analysis.

### SCD1 overexpression via adenovirus

The replication defective adenovirus Ad5/F35 was designed using the licensed Vector Builder software. The vector pAd5/F35[Exp]-mCherry-Cd144>mSCD1[NM_009127.4] contains the Cdh5 promoter to confer EC specificity, mCherry as a reporter to allow for visualization of targeted ECs, and the sequence of mouse *Sdc1* [NM_009127.4]. The adenovirus was generated by standard laboratory procedures in BSL2 biosafety cabinets using 293A cells. On the day of transfection, recombinant adenoviral plasmids were digested by the restriction enzyme PacI. The cells were transfected with Lipofectamine-DNA mix (Thermo Fisher Scientific) according to the manufacturer instructions. Lipofectamine-DNA mix was added to the culture flasks, which were returned to CO_2_ incubator at 37°C. The virus was left to infect the cells and then purified through a centrifugation process. Control adenovirus was (pAd5/F35[Exp]-mCherry-Cd144).

### Metabolomic Analysis

The metabolomic analyses in mouse plasma samples or endothelial cell lysates were performed at the West Coast Metabolomics Center at the University of California Davis, CA. Gas chromatography time-of-flight mass spectrometry (GC-TOF-MS) untargeted analysis identified 170 known metabolites and 290 unknown compounds. Metabolites were reported with retention index, quantification mass, and full mass spectra. Relative quantification was assessed by peak height. Targeted assays for selected fatty acid metabolites were performed for absolute quantification using stable isotope-labeled internal standards.

The data analysis was performed using MetaboAnalyst 5.0 (Research Resource Identifier RRID:SCR_015539). Before performing PCA, the concentrations of subjected metabolites were corrected using the Pareto scaling method to normalize the measurements of different metabolites on the close scale. The *x*-axis in the figures was depicted as the first principal component (PC1) representing the space with the largest variance in data, whereas the *y*-axis was depicted as the second principal component (PC2) representing the space with the second largest variance. The ovals are 95% inertia ellipses.

### Single Cell RNA Sequencing

The complete aorta was perfused with cold PBS, isolated and dissected to eliminate the surrounding periaortic fat. After mincing with scissors, a single cell suspension was obtained through enzymatic digestion with elastase (Worthington, Lakewood, NJ, USA) and liberase ((Roche, Basel, Switzerland) at 37°C. The preparation was passed through a 70 μm filter and treated with Red Blood Cell lysis buffer (Abcam, Waltham, MA, USA). Single cells were resuspended in PBS containing 0.04% BSA, counted and assessed for viability using Trypan Blue. Single cell RNA Sequencing was performed using the 10X Genomics Chromium platform at the UCLA Technology Center for Genomics and Bioinformatics. Raw sequencing data was demultiplexed and aligned using the Cell Ranger software from 10X Genomics, and we obtained the gene expression count matrix from Cell Ranger for downstream analyses. We sequenced ∼ 20,000 cells from the aorta. Quality control and data processing were performed in the Seurat package. PCA was used to reduce the number of features during the clustering stage. We used the modularity optimization algorithms implemented in Seurat to classify cells into clusters, and to identify the cell-type in each cluster using the cell-type-specific genes.

Seurat package was used to normalize count numbers by geometric mean and calculate the fold change and statistical significance, with *p* < 0.05 considered significant. Data were visualized using Python and Cytoscape.

### Single Cell RNA Data Analysis

The raw fastq scRNA reads were processed using Cell Ranger software (10x Genomics) and subjected to quality control with FastQC. The resulting output data for the three treatment groups were analyzed with the Scanpy package in Python. To generate an annotated data object, the data across groups were concatenated. Cells with gene counts above 4,000 or below 200 were filtered, as were those with a mitochondrial gene count exceeding 10%. Subsequently, we performed normalization, principal component analysis, variable gene extraction, non-linear dimensional reduction, and clustering to produce Uniform Manifold Approximation and Projection (UMAP) plots and identify cell clusters using cell specific marker genes. To visualize gene expressions between categories of related diet +/- exercise we created a circular heatmap. Circular heatmap features the log2 transformation of the fold changes, calculated as described above. Our heat maps were created using the heatmap.2 program in the ‘gplots’ package of R (http://cran.r-project.org). Circlize package R was used to generate circular heat map and hierarchical clustering. DESeq was used for normalization. All heat maps shown are row-normalized for presentation purposes.

### Statistics

All data are presented as mean ± standard error of the mean (SEM). Cell culture results consisted of 3 independent experiments. Graph Pad Prism software (GraphPad Software, La Jolla, CA) was used for statistical analysis. *t*-test or two-way ANOVA with multiple comparisons (genotype and presence or absence of exercise being the two dependent variables) and Tukey *post hoc* tests were performed to determine the statistical significance. *P* values < 0.05 were considered statistically significant.

### Study Approval

Animal experiments were performed in compliance with the Institutional Animal Care and Use Committee (IACUC) at the University of California Los Angeles (UCLA), under animal welfare assurance number A3196-01. The Animal Research Committee (ARC) reviewed all animal procedures performed at UCLA. The mice colony was housed in the facilities maintained by the UCLA Department of Laboratory Animal Medicine (DLAM). Mice obtained from external sources were allowed to acclimate for at least one week at UCLA facilities prior to the start of the experiments.

## AUTHOR CONTRIBUTIONS

SC, RL and TKH designed the study. SC, RL, HP, AMBM, JMC and SGR performed in vivo mouse studies and immunostaining. MR performed all ultrasound studies and CFD simulations. MR and SS performed aorta transcriptomics analysis with help from KV and JMC. SC, RL, and SS performed metabolomic analysis. SS performed bioinformatic network analysis. SC, JMC and SGR performed sample preparation for transcriptomic analysis. SS designed and prepared SCD1 overexpressing adenovirus. SC, RL, KIB and JJM performed *in vitro* experiments in HAEC. CDS and SR provided resources. SKP, TY, JAS, JJM, and STR provided intellectual feedback. SC, MR, SS and TKH wrote the manuscript. All authors discussed the results and participated in the article and figure preparation and editing.

## ACKNOWLEDGMENTS

The authors wish to express gratitude to Dr. Srinivasa T. Reddy (UCLA) and Dr. Alan M. Fogelman (UCLA) for providing the *Ldlr*^-/-^, *Scd1*^f^*^loxed/foxed^* mice, Dr. Curt D. Sigmund (Medical College of Wisconsin) for the EC-DN-PPARγ mice, Dr. Luisa Iruela-Arispe (Northwestern University) for *Cdh5*Cre mice, Dr. James G. Tidball (UCLA) for providing mouse exercise wheels and critical feedback and Dr. Alison Marsden (Stanford University) for the SimVascular Code for Computational Fluid Dynamics. The authors are grateful for Dr. Ronald McGregor’s assistance with the confocal analysis and Ms. Vienna Benavides for assistance with *in vitro* experiments. This work was supported by funds from the Department of Veterans Affairs VA Merit Award I01 BX004356 (T.K.H) and National Institutes of Health NIH R35HL144807 (C.D.S), R01HL118650 (T.K.H), R01HL111437 (T.K.H) and R01HL149808 (T.K.H).

